## Supplemental Information PDF for "Variation in anthelmintic responses are driven by genetic differences among diverse *C. elegans* wild strains"

### DESCRIPTION OF ADDITIONAL SUPPLEMENTARY TABLES

**Supplemental Table 1.** Strain-specific $EC_{10}$ estimates for each anthelmintic (µM units).

**Supplemental Table 2.** Strain-specific slope estimates for each anthelmintic.

**Supplementary Table 3.** Relative potency estimates in pairwise comparisons of $EC_{10}$ estimates among all strains for each anthelmintic.

**Supplementary Table 4.** Relative potency estimates in pairwise comparisons of slope estimates among all strains for each anthelmintic.

**Supplementary Table 5.** Anthelmintic drug stock solution preparation details.

### SUPPLEMENTARY FIGURES

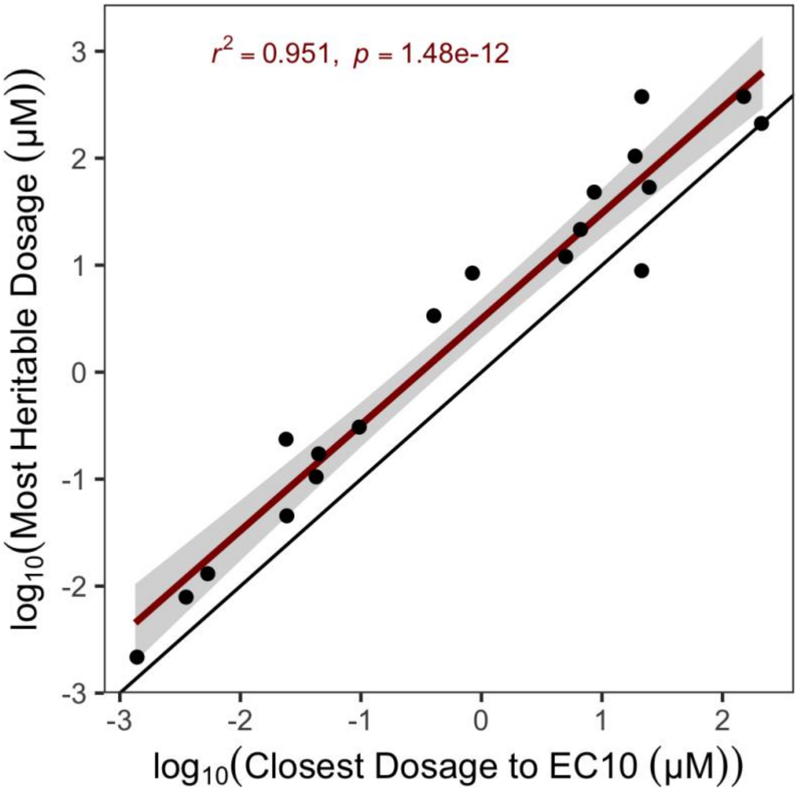


**Supplemental Figure 1. $EC_{10}$ estimates from genetically diverse strains predict exposures with heritable responses.** The log-transformed exposure that elicited the most heritable response to each anthelmintic (y-axis) is plotted against the log-transformed exposure of that same anthelmintic nearest to the inferred $EC_{10}$ from the dose-response assessment. The exposure closest to the $EC_{10}$ across all anthelmintics exhibited significant explanatory power to determine the exposure that elicited heritable phenotypic variation.

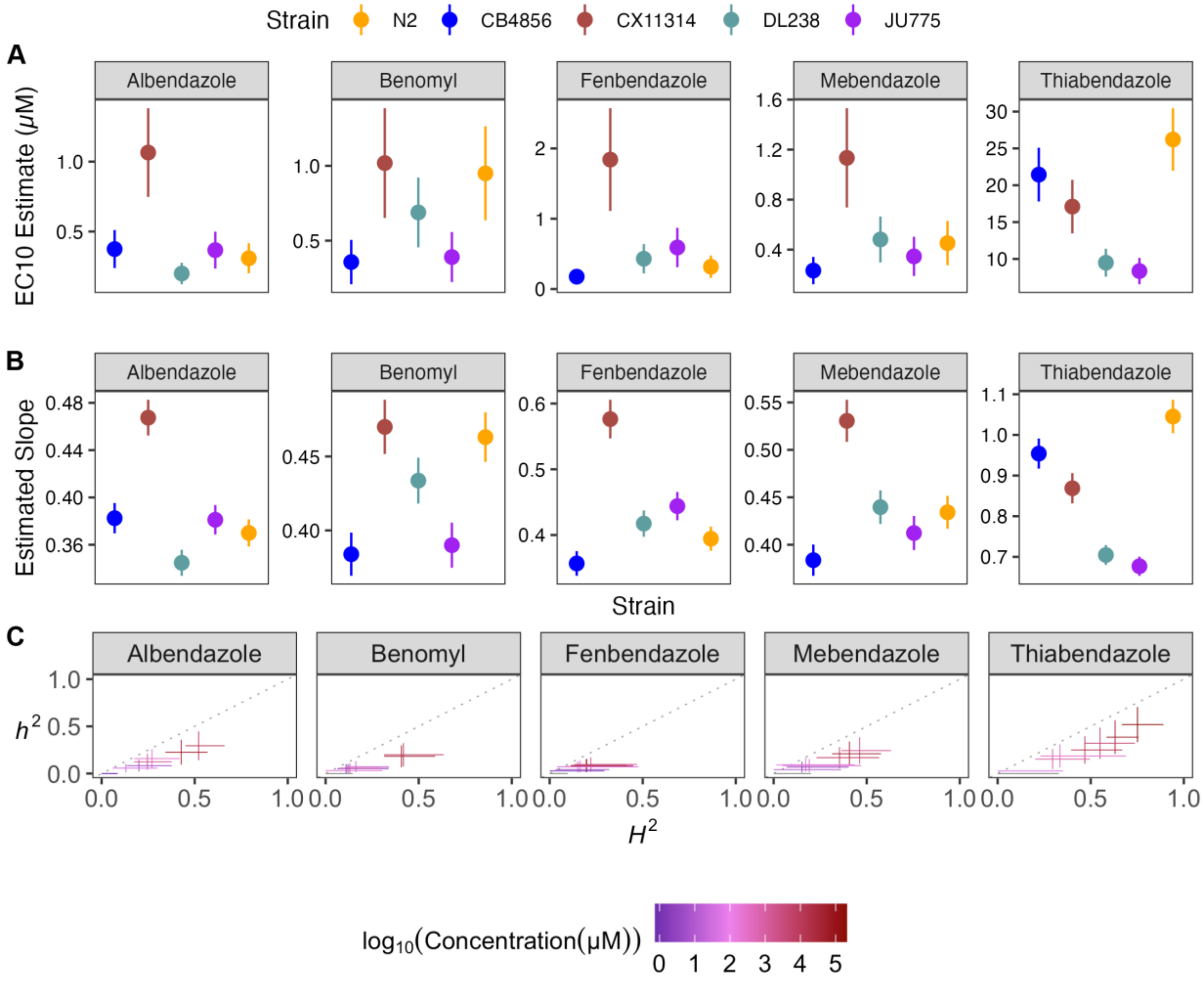


**Supplementary Figure 2. Variation in benzimidazole (BZ) $EC_{10}$ dose-response and slope estimates without MY16 demonstrate how small genetic effects have variable responses across strains. A)** Strain-specific $EC_{10}$ estimates (*e*) for each benzimidazole are displayed for each strain. Standard errors for each strain- and anthelmintic-specific $EC_{10}$ estimates are shown. **B)** Strain-specific slope estimates (*b*) for each benzimidazole are displayed for each strain. Standard errors for each strain- and anthelmintic-specific slope estimate are indicated by the line extending vertically from each point. **C)** The broad-sense (x-axis) and narrow-sense heritability (y-axis) of normalized animal length measurements were calculated for each concentration of each benzimidazole (*Methods*; *Broad-sense and narrow-sense heritability calculations*). The color of each cross corresponds to the log-transformed dose for which those calculations were performed. The horizontal line of the cross corresponds to the confidence interval of the broad-sense heritability estimate obtained by bootstrapping, and the vertical line of the cross corresponds to the standard error of the narrow-sense heritability estimate.

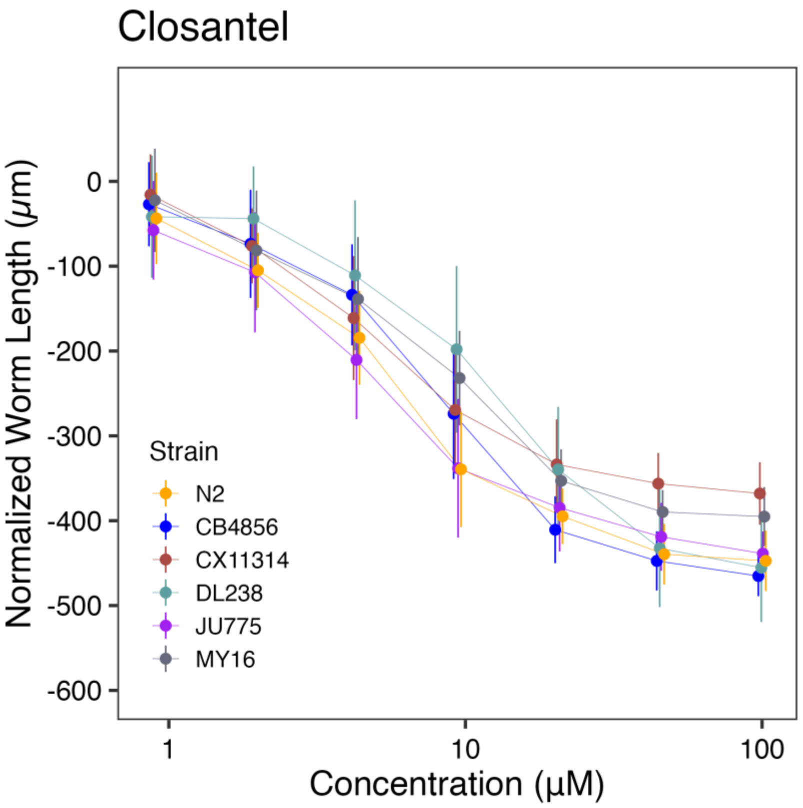
Closantel
Normalized Worm Length (μm)
0
-100
-200
-300
-400
-500
-600
Strain
N2
CB4856
CX11314
DL238
JU775
MY16
1
10
100
Concentration (μM)

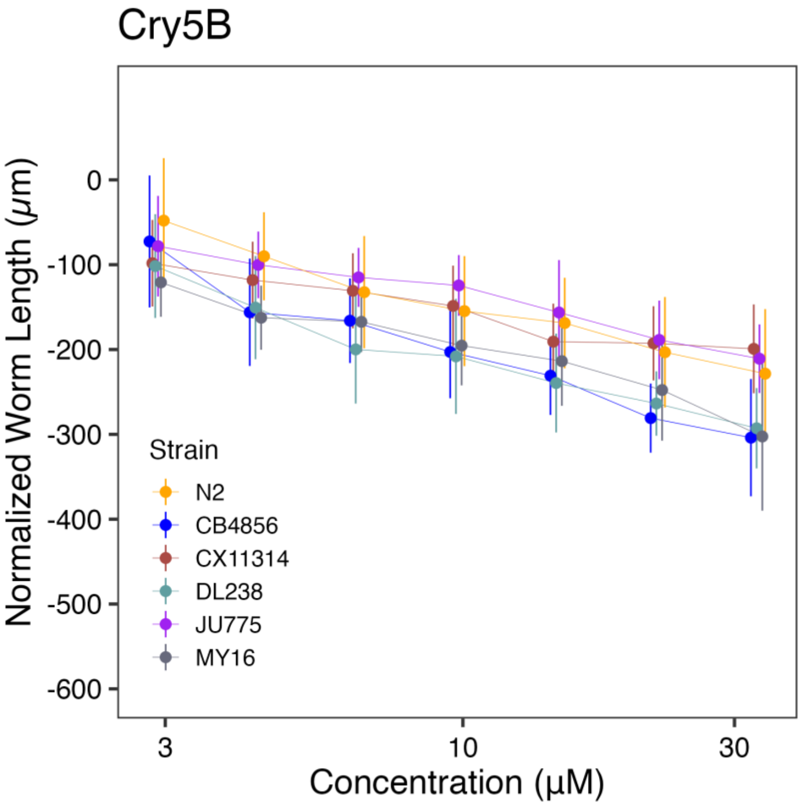
Cry5B
Normalized Worm Length (μm)
0
-100
-200
-300
-400
-500
-600
Strain
N2
CB4856
CX11314
DL238
JU775
MY16
3
10
30
Concentration (μM)

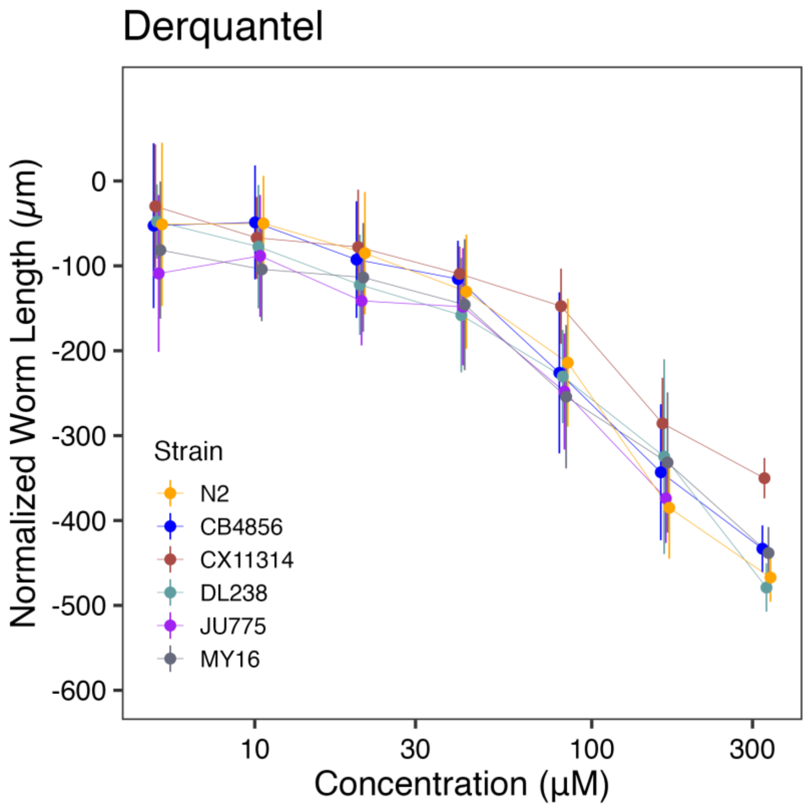

Derquantel
Normalized Worm Length (μm)
0
-100
-200
-300
-400
-500
-600
Strain
N2
CB4856
CX11314
DL238
JU775
MY16
10
30
100
300
Concentration (μM)


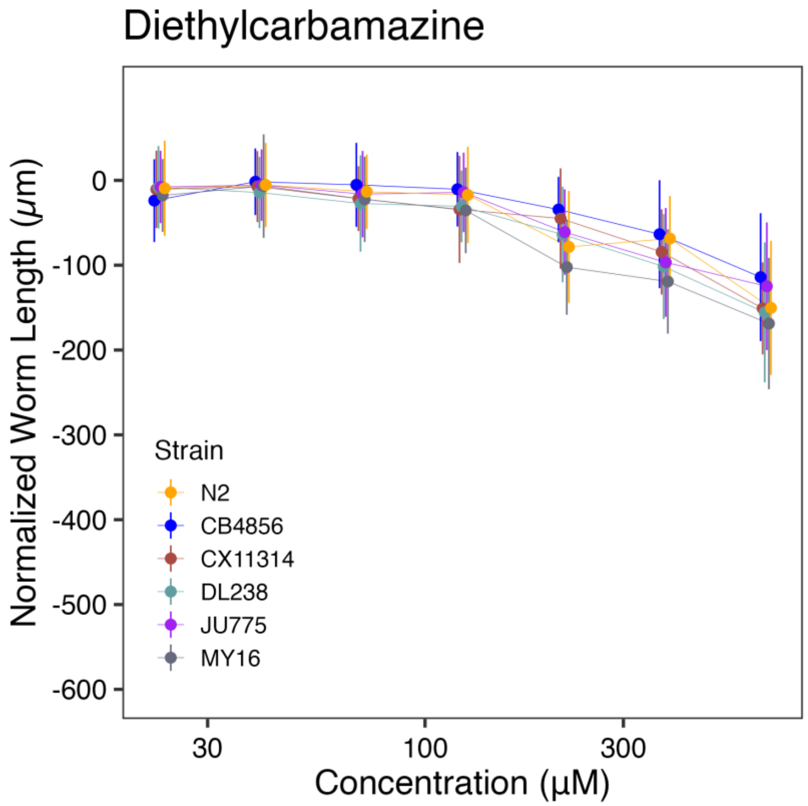

Diethylcarbamazine
Normalized Worm Length (μm)
0
-100
-200
-300
-400
-500
-600
Strain
N2
CB4856
CX11314
DL238
JU775
MY16
30
100
300
Concentration (μM)

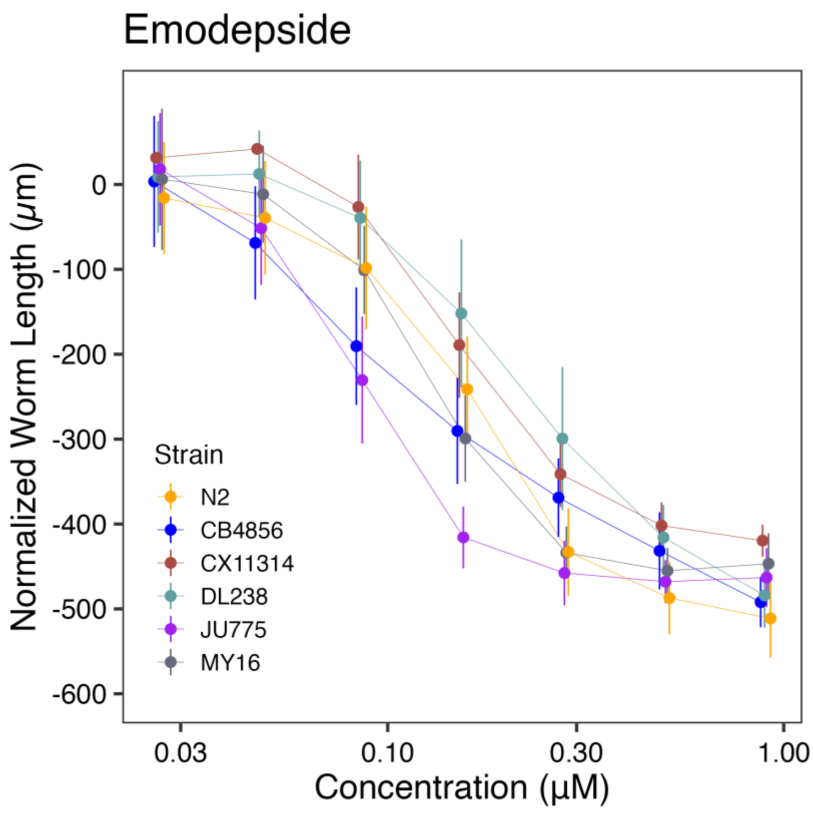
Emodepside
Normalized Worm Length (µm)
0
-100
-200
-300
-400
-500
-600
Strain
N2
CB4856
CX11314
DL238
JU775
MY16
0.03
0.10
0.30
1.00
Concentration (µM)

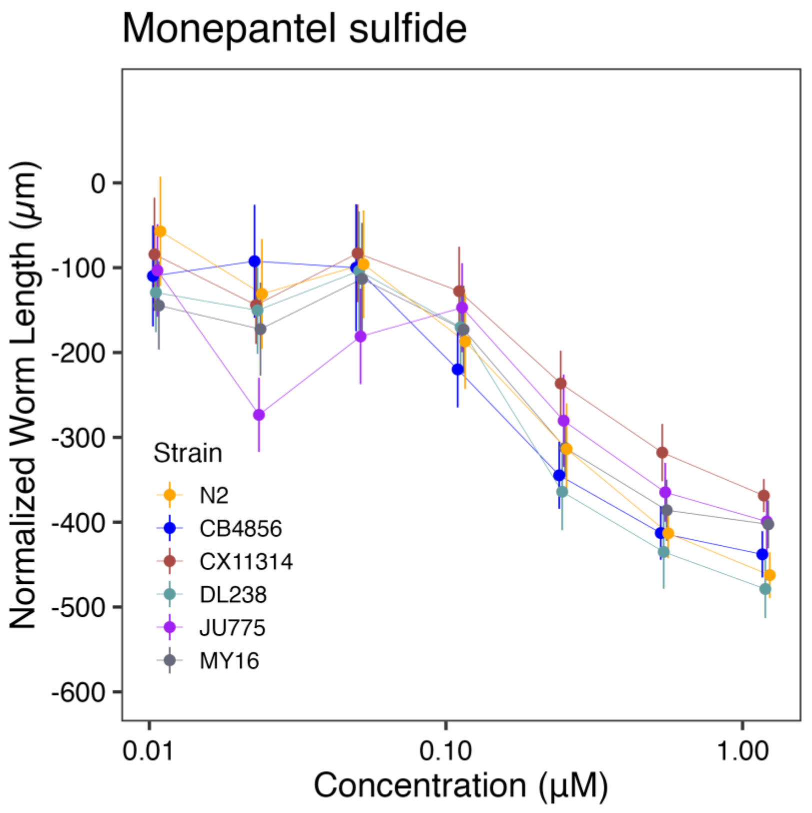
Monepantel sulfide
Normalized Worm Length (µm)
0
-100
-200
-300
-400
-500
-600
Strain
N2
CB4856
CX11314
DL238
JU775
MY16
0.01
0.10
1.00
Concentration (µM)

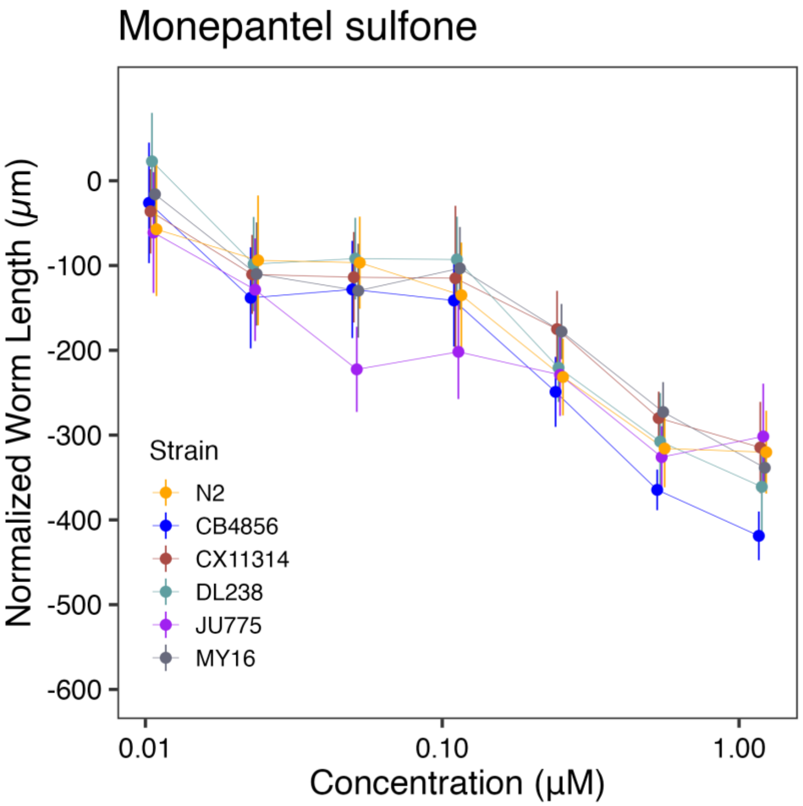
Monepantel sulfone
Normalized Worm Length (μm)
0
-100
-200
-300
-400
-500
-600
0.01
0.10
1.00
Concentration (μM)
Strain
N2
CB4856
CX11314
DL238
JU775
MY16

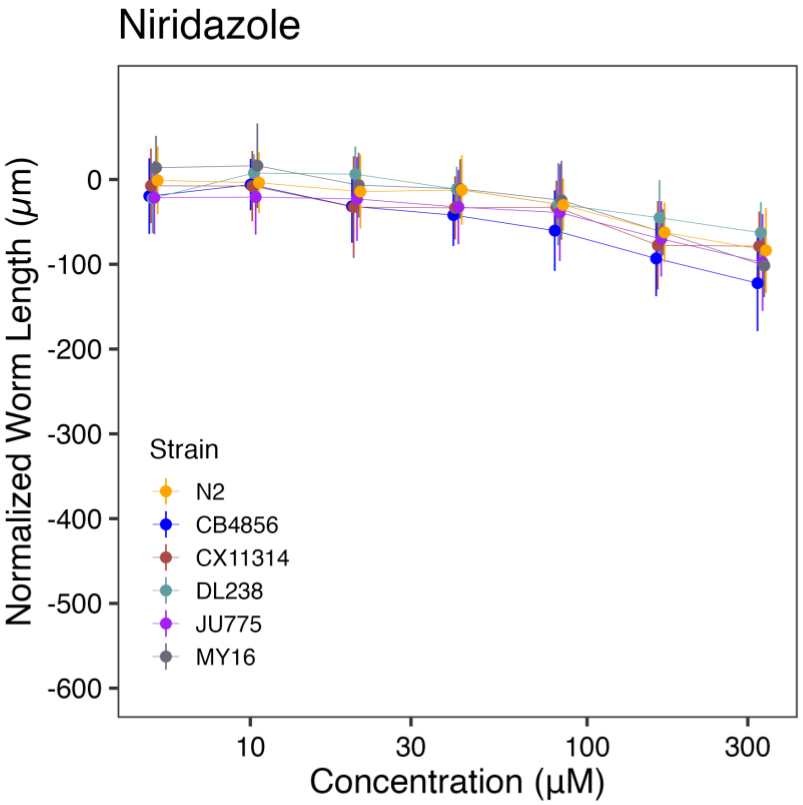
Niridazole
Normalized Worm Length (μm)
0
-100
-200
-300
-400
-500
-600
10
30
100
300
Concentration (μM)
Strain
N2
CB4856
CX11314
DL238
JU775
MY16

Oxamniquine

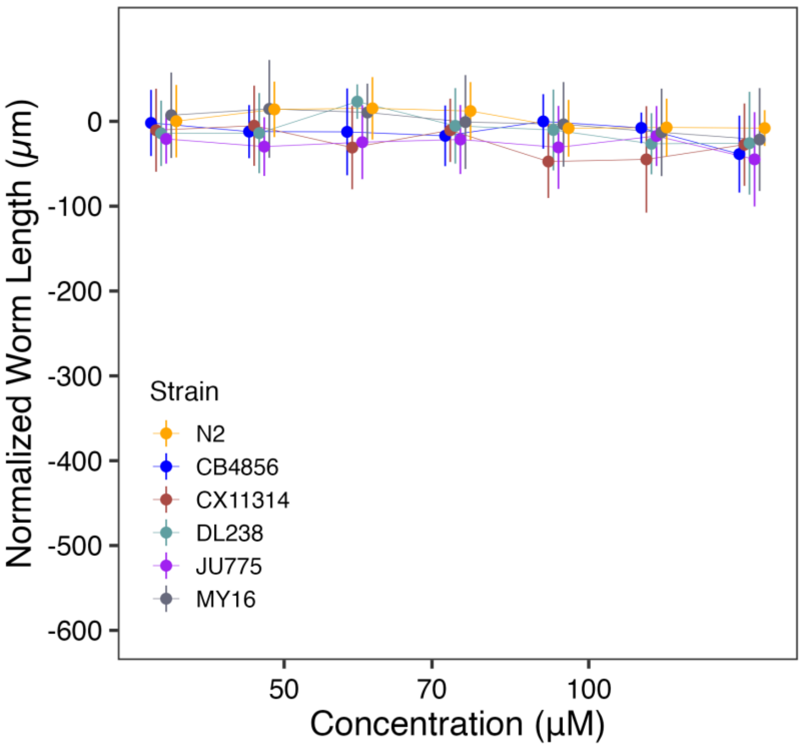


Piperazine

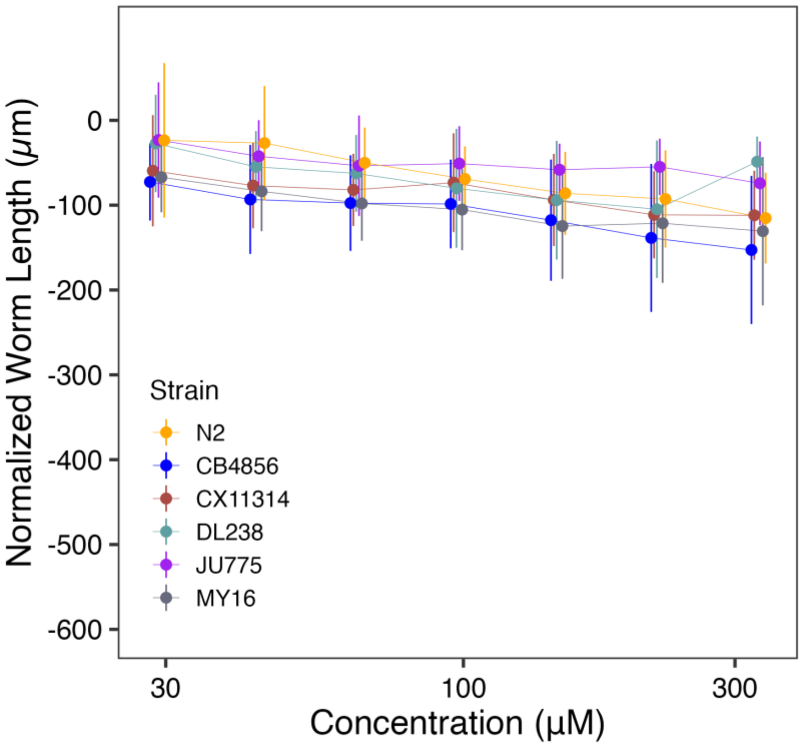

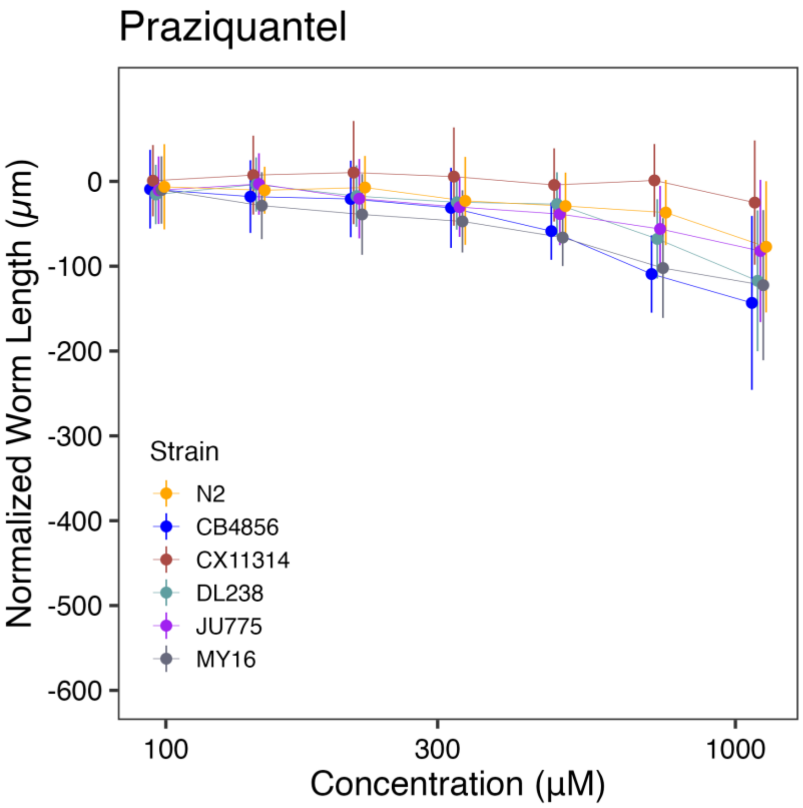


**Supplementary Figures 3 - 13: Dose-response curves for anthelmintic drugs and nematicides not in the three main anthelmintic drug classes.** Normalized animal lengths (y-axis) are plotted for each strain as a function of the dose of anthelmintic supplied in the high-throughput microscopy assay (x-axis). Strains are denoted by color. Lines extending from points represent the standard deviations from the mean responses. Statistical normalization of animal lengths is described in *Methods.*

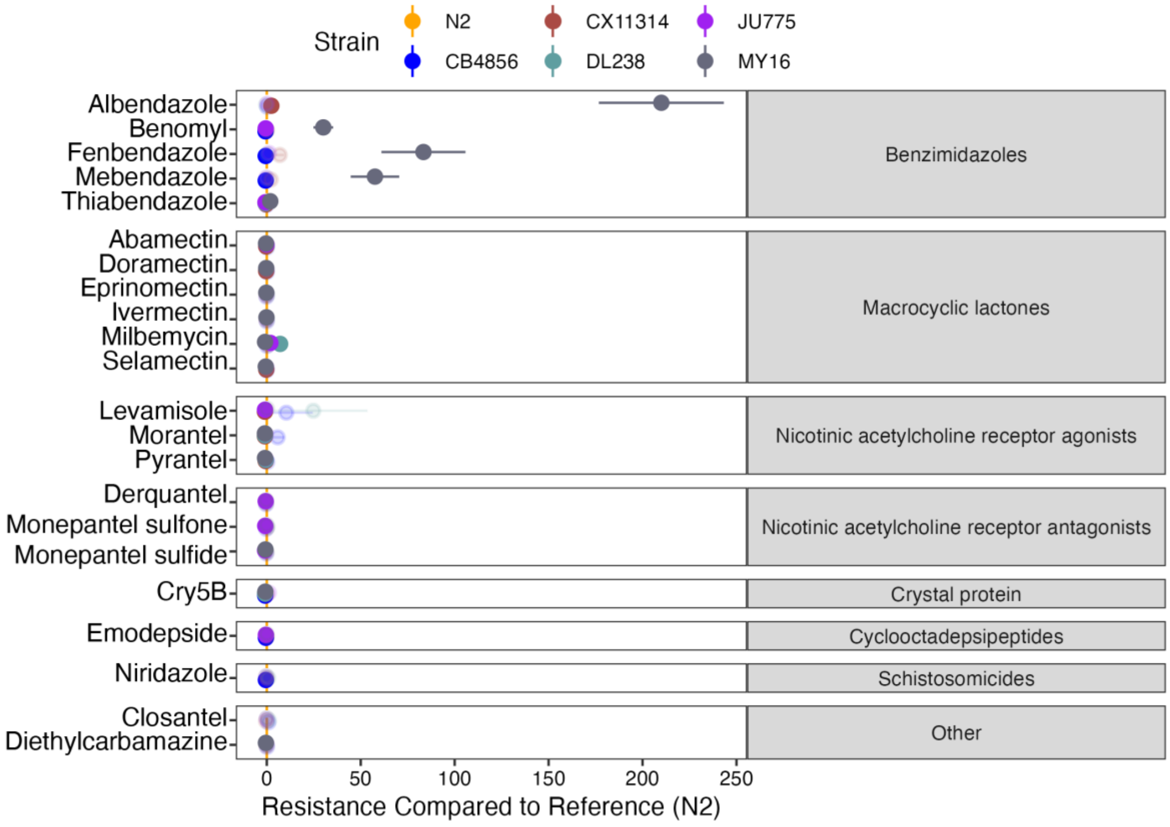


**Supplementary Figure 14. Variation in $EC_{10}$ estimates can be explained by genetic variation across strains.** For each anthelmintic, the relative potency of that anthelmintic against each strain compared to the N2 strain is shown. Solid points denote strains with significantly different relative resistance to that anthelmintic (Student's t-test and subsequent Bonferroni correction with a $p_{adj} < 0.05$), and faded points denote strains not significantly different than the N2 strain. The broad category to which each anthelmintic belongs is denoted by the strip label for each facet. Anthelmintic drugs with undefined $EC_{10}$ estimates (estimates greater than the maximum dose to which animals were exposed) are not shown.

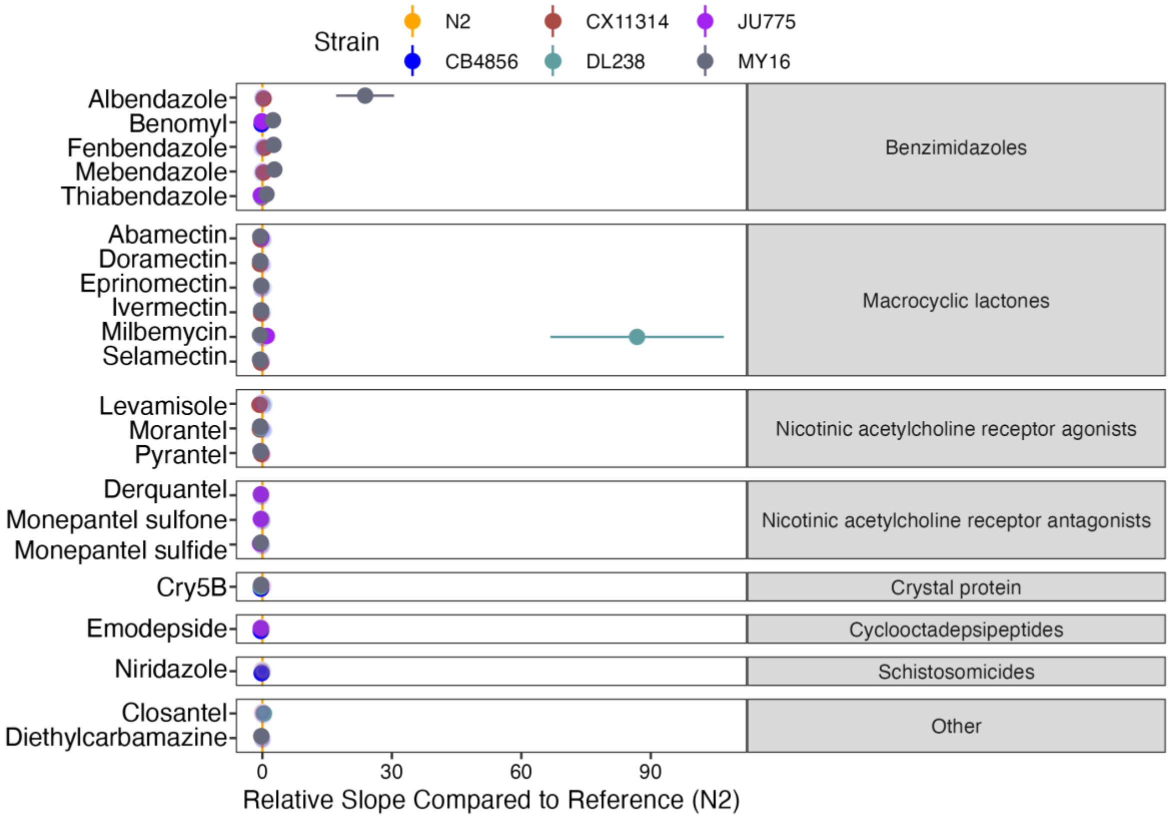


**Supplementary Figure 15. Variation in dose-response slope estimates can be explained by genetic differences among strains.** For each anthelmintic, the relative steepness of the dose-response slope inferred for that strain compared to the N2 strain is shown. Solid points denote strains with significantly different dose-response slopes (Student's t-test and subsequent Bonferroni correction with a $p_{adj} < 0.05$), and faded points denote strains without significantly different slopes than the N2 strain. The broad category to which each anthelmintic belongs is denoted by the strip label for each facet. Anthelmintic drugs with undefined slope estimates are not shown.

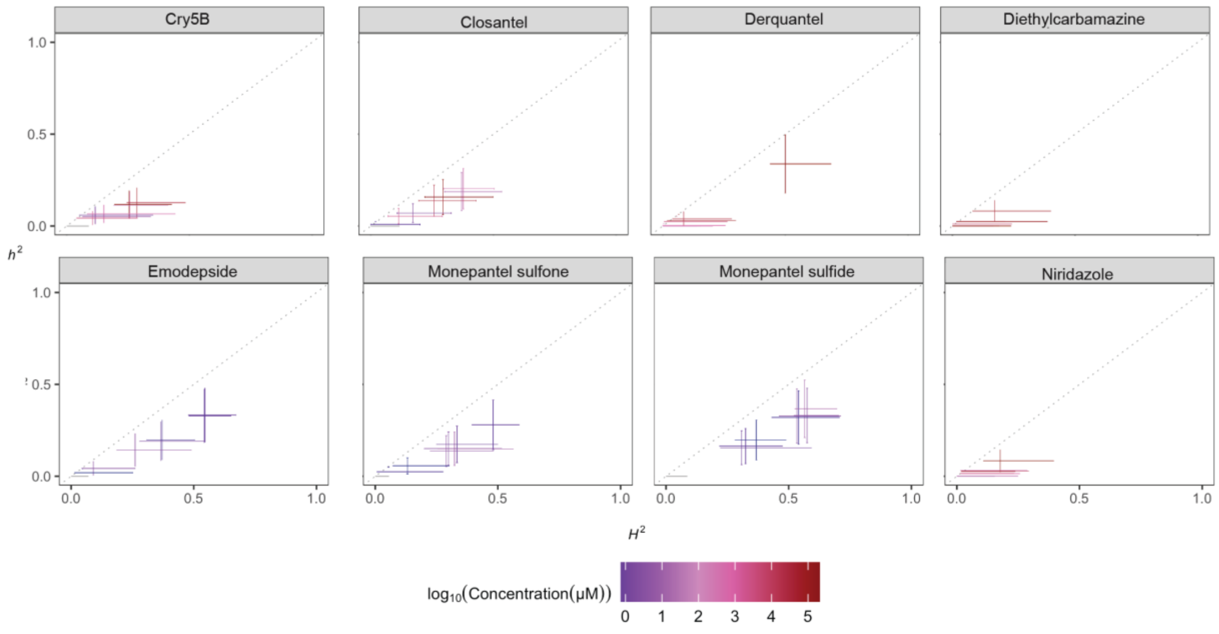


**Supplementary Figures 16: Heritability plots for anthelmintic drugs and nematicides not in the three main anthelmintic drug classes.** The broad-sense (x-axis) and narrow-sense heritability (y-axis) of normalized animal length measurements were calculated for each concentration of each nicotinic acetylcholine receptor agonist (*Methods*; *Broad-sense and narrow-sense heritability calculations*). The color of each cross corresponds to the log-transformed dose for which those calculations were performed. The horizontal line of the cross corresponds to the confidence interval of the broad-sense heritability estimate obtained by bootstrapping, and the vertical line of the cross corresponds to the standard error of the narrow-sense heritability estimate. Heritability could not be calculated for anthelmintics without an $EC_{10}$.

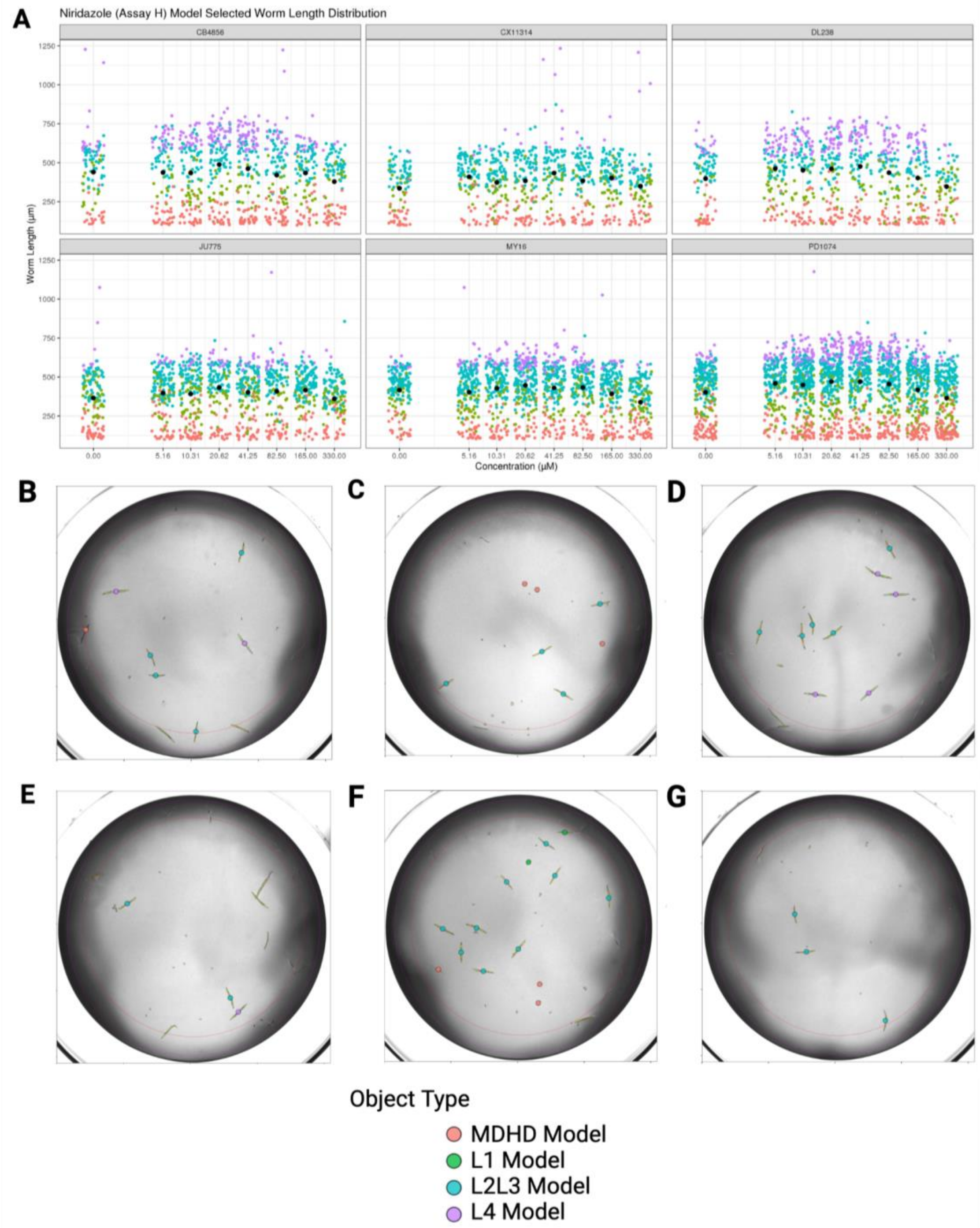


**Supplementary Figure 17. Problematic assays and well features for downstream analysis. A)** Distribution of animals by length (y-axis) is plotted for each strain by dose (x-axis) for niridazole in Assay H. Each data point is a single observed object. Data points are colored by worm model: L4 larval stage (purple), L2 and L3 larval stages (turquoise), L1 larval stage (green), and MDHD (pink). The black data point denotes the average worm length by dose. Well images obtained in Assay H for niridazole dose-response experiments are representative of problematic control well features caused by the presence of small animals categorized by the L1 model and L2L3 for each strain: **B)** CB4856, **C)** CX11314, **D)** DL238, **E)** JU775, **F)** MY16, and **G)** N2.

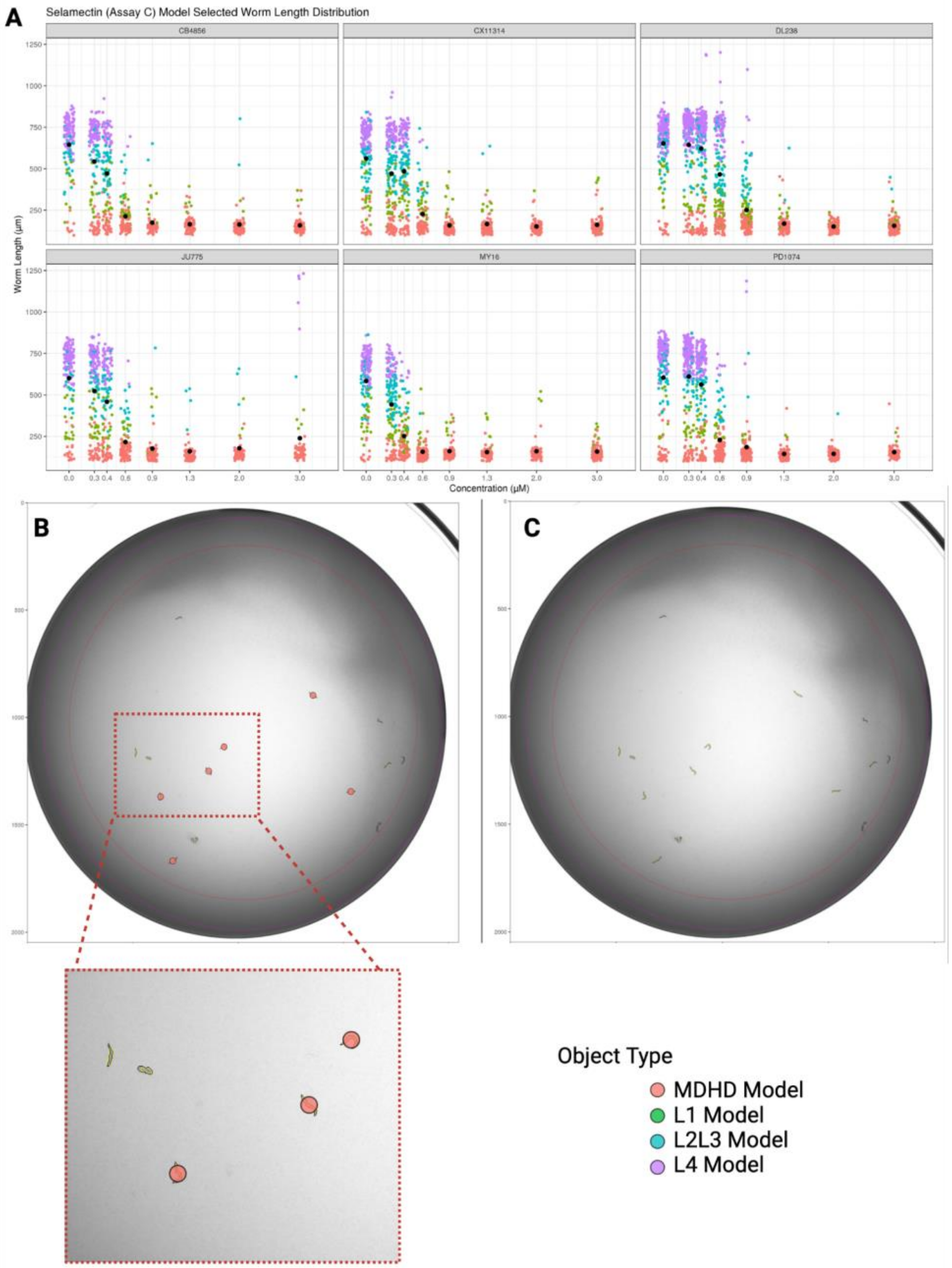


**Supplementary Figure 18. Selamectin benefits from retaining *Worm_Length* > 30 objects and the MDHD model. A)** Distribution of animals by length (y-axis) is plotted for each strain by dose (x-axis) for selamectin in Assay C. Each data point is a single observed object. Data points are colored by worm model: L4 larval stage (purple), L2 and L3 larval stages (turquoise), L1 larval stage (green), and MDHD (pink). Average worm length by dose is denoted by the black datapoint. Selamectin high dose wells are shown for both the **B)** *Worm_Length* > 30-pixel filter and **C)** *Worm_Length* > 50-pixel filter. The inset shows small animals recognized by the MDHD model. Pink overlays indicate an MDHD model selected object. Animals with no overlay are filtered out during the cleaning steps**.** See *Methods* for details on data cleaning metrics.
